# Fairy: fast approximate coverage for multi-sample metagenomic binning

**DOI:** 10.1101/2024.04.23.590803

**Authors:** Jim Shaw, Yun William Yu

## Abstract

**Background:** Metagenomic binning, the clustering of assembled contigs that belong to the same genome, is a crucial step for recovering metagenome-assembled genomes (MAGs). Contigs are linked by exploiting consistent read coverage patterns across a genome. Using coverage from multiple samples leads to higher-quality MAGs; however, standard pipelines require all-to-all read alignments for multiple samples to compute coverage, becoming a key computational bottleneck.

**Results:** We present fairy (https://github.com/bluenote-1577/fairy), an *approximate* coverage calculation method for metagenomic binning. Fairy is a fast k-mer-based alignment-free method. For multi-sample binning, fairy can be ***>* 250*×*** faster than read alignment and accurate enough for binning. Fairy is compatible with several existing binners on host and non-host-associated datasets. Using MetaBAT2, fairy recovers **98.5%** of MAGs with ***>* 50%** completeness and ***<* 5%** incompleteness relative to alignment with BWA. Notably, multi-sample binning with fairy is *always* better than single-sample binning using BWA (***>* 1.5*×*** more ***>* 50%** complete MAGs on average) while still being faster. For a public sediment metagenome project, we demonstrate that multisample binning recovers higher quality Asgard archaea MAGs than single-sample binning and that fairy’s results are indistinguishable from read alignment.

**Conclusions:** Fairy is a new tool for approximately and quickly calculating multi-sample coverage for binning, resolving a longstanding computational bottleneck for metagenomics.

## 1 Background

Direct shotgun sequencing of microbiomes has allowed for the recovery of metagenome-assembled genomes (MAGs), unlocking unprecedented insights into the ecology of even unculturable organisms [1]. The computational process of generating MAGs first requires *assembling* the sequenced reads into contiguous sequences called *contigs*. After assembly, contigs are grouped into MAGs by *binning* through either automated algorithms [2–5] or manual curation [6].

Binning is done by leveraging consistent information across an *entire genome*. Such genomic signatures [7] include k-mer frequencies (e.g. tetranucleotide frequencies) and sequencing coverage, which are commonly used in binning algorithms. Combining coverage information from *multiple samples* is an effective way of increasing the resolving power of binning algorithms [8]. It has been shown that multi-sample coverage is vastly superior to single-sample coverage for binning and produces better MAGs that even quality-control software such as CheckM [9] may not be able to detect [10].

Coverage calculation is usually done by aligning reads back to contigs (e.g. using BWA [11] or BowTie2 [12]). For a project with *n* samples and *n* resulting assemblies, computing multi-sample coverage naively requires aligning each sample to each assembly, resulting in *n*^2^ read-alignment runs. This quadratic scaling becomes prohibitively long when the number of samples is large. Co-assembly, where all reads are assembled to give one set of contigs, is a potential solution, but co-assembly can be memory intensive and collapse similar strains [13]. Another method is split-binning [4], where all contigs from a set of assemblies are concatenated together and then aligned to. This is faster but still relatively time and memory-intensive. Thus, many large-scale studies still do single-sample binning [14].

If only coverage is needed, read alignment is computationally wasteful because the exact base alignments are not needed. Alignment-free methods are faster and provide an intriguing alternative. For example, pseudoalignment [15, 16] has been used for binning coverage [17, 18]. Additionally, direct k-mer counts have also been used to separate strains in de Bruijn graphs [19] and applied to multi-sample coverage for binning [20]. However, we are not aware of any customized tools for this task with detailed benchmarks.

### 1.1 Our contributions

In this paper, we present a much faster, k-mer-based alignment-free method of computing multi-sample coverage for metagenomic binning. Our method, *fairy*, is built on top of our metagenomic profiler *sylph* [21], but fairy is specifically adapted for metagenomic binning of contigs. The key computational insight of fairy is that indexing the *reads* with a sparse set of k-mers is much more efficient than indexing the *genomes*. This is due to the redundancy of k-mers within the reads, allowing fairy to be 2-3 orders of magnitude faster than alignment. A variant of this technique was previously employed for compressively accelerated all-mapping, but that task still required base-level alignment, whereas fairy eschews that step [22]. While fairy’s coverages are approximate, we show that approximate coverages are good enough for multi-sample binning.

## 2 Methods

### 2.1 Repurposing a fast metagenomic profiler for approximate coverage computation

Fairy’s codebase was forked and independently developed from sylph, a metagenomic profiler we developed [21]. However, fairy is extended and repurposed for metagenomic binning. Fairy and sylph are algorithmically similar, but they differ in three ways:

1. **Default parameter choices**. We will make the differences explicit below.
2. **User interface**. This includes command line options, inputs, and output formats – fairy is a *coverage calculator* rather than a *metagenomic profiler*.
3. **A coverage variance computation step**. This is useful for some binners such as MetaBAT2 [2].

Fairy’s output is the same as the commonly used `jgi_summarize_bam_contig_depths` script from MetaBAT2 and can be toggled to be compatible with MaxBin2 as well. We recapitulate fairy’s key algorithmic steps below, also shown in **Fig. 1A**. We outline technical details pertaining to sylph’s algorithm in Appendix A.

**Fig. 1.**
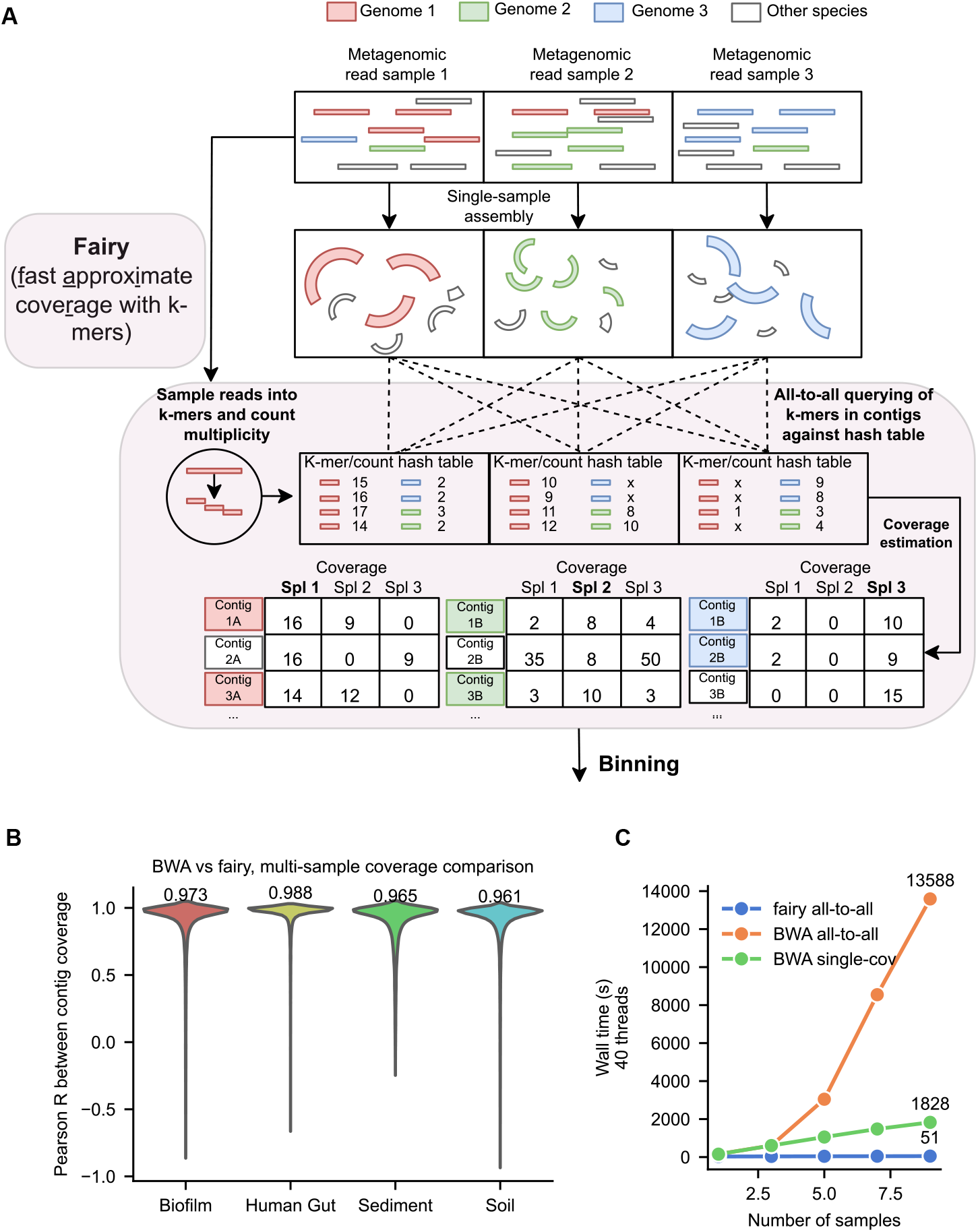
Fast approximate k-mer coverage estimates for multi-sample metagenomic binning. **A**. Outline of fairy’s k-mer-based algorithm. Fairy’s processing steps are outlined in light red. Fairy indexes (or *sketches*) the reads into subsampled k-mer-to-count hash tables. K-mers from contigs are then queried against the hash tables to estimate coverage. Finally, fairy’s output is used for binning and is compatible with several binners (e.g. MetaBAT2, MaxBin2). **B**. Pearson R values between fairy and BWA’s multi-sample coverages for contigs in an arbitrary assembly from the dataset. Median values are shown above the plots. **C**. Wall time with 40 threads for fairy vs BWA on 1, 3, 5, 7, and 9 gut samples in all-to-all mode (multi-sample) and single-sample mode.

#### 2.1.1 Sparse sketching of k-mers

Fairy sparsely samples k-mers from reads and assemblies using the FracMinHash [23] method to sample approximately 1*/*50 k-mers (whereas sylph defaults to a 1*/*200 sampling rate). For each metagenomic sample, the k-mers in the reads and their associated multiplicities within the sample are stored in a hash table. The hash table indices (one for each sample) are written to disk and loaded as needed, whereas the assemblies’ k-mers are kept in memory.

#### 2.1.2 Contig querying and containment

For an assembly, every contig’s sampled k-mers are queried against every metagenomic sample’s hash table. For a contig, fairy requires 8 k-mers at minimum to be contained in the sample (whereas sylph requires 50) to proceed with coverage calculation. In addition to the minimum k-mer threshold, fairy calculates a *containment ANI* in the same manner as sylph (Appendix A). Containment ANI measures the average nucleotide identity of the contig to all sequences in the metagenome. If the containment ANI is *<* 95%, fairy assumes the contig is not present in the sample at species-level, and thus assigns a coverage of 0.

We choose 95% as a presence-absence threshold by default, corresponding to species-level ANI thresholds used in practice [24]. Strain-level MAG binning from strain-resolved assemblies (e.g. with PacBio HiFi reads [25]) may require higher ANI thresholds, but higher thresholds would reduce sensitivity for multi-sample binning. In the results section, we discuss some issues with the 95% threshold for PacBio HiFi MAG binning.

### 2.2 Coverage calculation from k-mers

For a contig passing the containment ANI threshold, the coverage is estimated in three different ways. Let *M* be the median k-mer multiplicity of the contig’s k-mers within the sample. We calculate the coverage exactly as is done with sylph’s “effective coverage” estimator [21], briefly restated below (see Appendix A for more information). Fairy’s output coverage is:

- If *M* ≤ 3: a statistical estimator using Poisson coverage assumptions.
- If 4≤ *M* ≤ 15: a robust mean of k-mer counts, trimming off large k-mer counts.
- If *M >* 15: median of the k-mer counts.

#### 2.2.1 Coverage variance calculation

The variance is calculated as the sample variance of the contig’s non-zero k-mer multiplicities in the sample with the 10 and 90 percentile k-mer multiplicities trimmed to remove long-tailed k-mer coverage outliers (e.g. due to mobile elements).

### 2.3 Why fairy is fast for multi-sample computation

Fairy’s speed comes from the fact that each sample’s reads are only processed once. Processing reads into hash tables turns out to be far more computationally costly than querying a contig’s k-mers against the hash table. Thus, for *n* samples, fairy only requires *n* costly read-processing steps and *n*^2^ fast query procedures. This makes fairy scale well for multi-sample binning compared to read alignment.

### 2.4 Benchmarking procedure

We benchmark on real datasets with multiple samples [26–32] and subsetted the samples when the number was too large; exact accessions are available in **Supplementary Table 2**. We opted to benchmark on real data as opposed to synthetic data to show that fairy has good performance on realistic datasets. It has been found that CheckM’s relative performance is strongly correlated with ground-truth performance on simulated data [33]. In particular, we primarily care about relative results rather than true contamination/completeness.

#### 2.4.1 Assembly and MAG generation

For generating assemblies and MAGs using read alignment for binning, our procedure was as follows:

1. **Short-read metagenomes**: we generated assemblies using ATLAS [34] v2.18.1 with default settings. We mapped short-reads using BWA and used CoverM [35] with the `-m metabat` option, which is identical to using the `jgi_summarize_bam_contig_depths` script from MetaBAT2 with default settings.
2. **Nanopore long-read metagenomes**: we generated assemblies using metaFlye [36] with default settings. We mapped reads using minimap2 [37] and used the `jgi_summarize_bam_contig_depths` script with minimum mapping quality 5 and alignment identity 80% (as opposed to default 97%) due to lower accuracy of nanopore reads.
3. **PacBio HiFi metagenomes**: we used MetaMDBG [38] for assembly and minimap2 with `jgi_summarize_bam_contig_depths` (default settings) for coverage calculation.

For all above data types, fairy v0.5.1 was then used with default parameters on the resulting assemblies and reads to generate another set of coverage profiles. Fairy’s coverage profiles were used to generate another set of MAGs that we compared to the MAGs generated from alignment-based coverages.

### 2.4.2 Binning and evaluation

For short-read assemblies, binning was conducted with MetaBAT2 v2.15 [2], MaxBin2 v2.2.7 [3], MetaBinner v1.4.4 [5] and VAMB v3.0.9 [4]. Notably, we used an older version of VAMB because the newer version required BAM inputs, which fairy can not generate. For long-read assemblies, we only used MetaBAT2, as it seemed to be more commonly used for existing long-read pipelines [25, 31, 38]. All binners were run with standard settings, except using a minimum contig length of 1500 when possible to emulate existing pipelines [10]. Some binners can only be run with a BAM file as opposed to a custom coverage profile, so we could not benchmark them [39, 40]. Finally, we use CheckM2 [9] to evaluate the contamination and completeness of the resulting bins.

## 3 Results

### 3.1 Fairy is concordant with read-alignment coverage while being *>* 100 times faster

For four short-read datasets (**Fig. 1B**), we took one arbitrary assembly and examined all present contigs. For *n* samples, each contig has *n* coverage values. We calculated the Pearson correlation between the *n* coverages that fairy output and BWA’s *n* coverages. The median Pearson R value was *>* 0.96 for all datasets, indicating good concordance. Sediment and soil metagenomes are more complex than gut metagenomes, likely explaining the lower concordance between BWA and fairy on these datasets (0.988 for gut vs 0.965 for sediment and 0.961 for soil).

Fairy is more than 250× faster than BWA for multi-sample coverage (**Fig. 1C**) when using just 9 of the short-read human gut metagenomes. While fairy technically requires a quadratic number of coverage calculations, much of the processing is in the linear-time indexing step. For read-alignment, the quadratic time cost of all-to-all alignment is clear for even a small number of samples.

#### 3.1.1 Multi-sample coverage calculation quickly becomes a bottleneck, but not for fairy

For the soil dataset (10 samples) with 40 cores, assembly with SPAdes took approximately 15 hours, while read alignment took more than 40 hours. Theoretically, if we used 100 samples instead of 10, assembly would take ≈15 × 10 = 150 hours and finish within a week, whereas all-to-all alignment would take 40× 10^2^ = 4000 hours (around 167 days), which is not feasible. On this dataset, fairy took 9 minutes for indexing and 7 minutes for querying; most of the querying time was spent on disk I/O, and we did not even use an SSD. In the worst case, 100 samples would still take less than a day for fairy.

For memory, fairy scales with the size and complexity of the metagenome. Processing the 10 aforementioned soil samples in parallel took 40 GB of RAM (4 GB per sample), which is still relatively small compared to assembly, and if memory is a constraint, fewer samples can be processed in parallel.

### 3.2 Multi-sample binning with fairy is better than single-sample binning with alignment

We compared fairy’s binning results with BWA and minimap2 using MetaBAT2 as the binner (**Fig. 2**). For every short-read dataset, multi-sample binning outperforms single-sample binning. This is especially prominent for complex sediment or soil metagenomes. We also note that single-sample binning contains contamination which may not be detectable by CheckM [10]. The long-read sludge metagenome is an exception, with minimap2 performing worse for multi-sample binning. We speculate that this is due to parameters (i.e. for coverage and binning) not being tuned as carefully for long reads. Compared to short-read binning, optimal parameter choices have not been explored as much for long-read binning – we do not claim our choice of alignment-based coverage parameters is optimal.

**Fig. 2.**
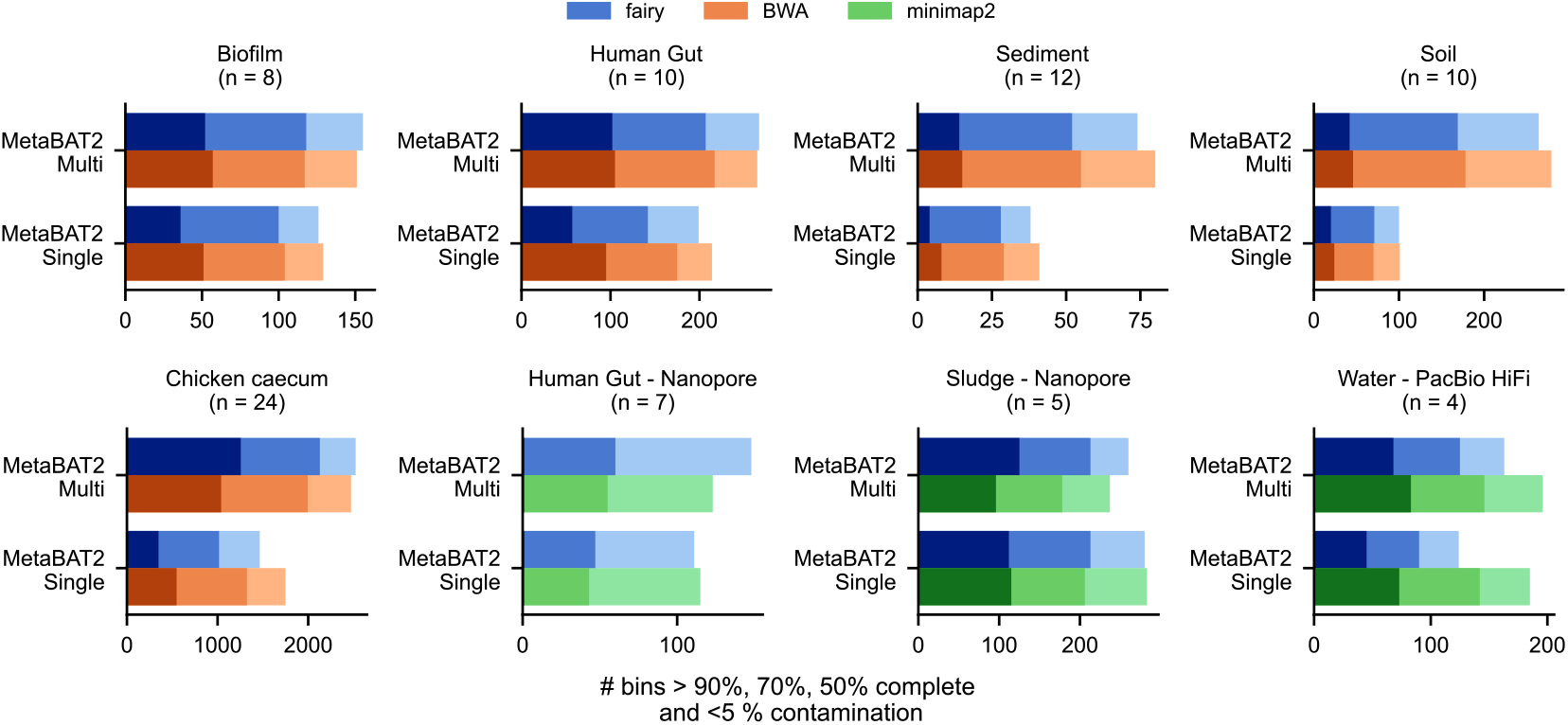
Binning results over multiple datasets and binners. Fairy’s concordance with BWA and minimap2 for multi-sample and single-sample coverage binning using MetaBAT2. minimap2 was used with long-read datasets (PacBio HiFi or nanopore) whereas BWA was used with short-read datasets. Sample accessions are available in Supplementary Table 2.

### 3.3 Fairy recovers comparable bins to alignment for multi-sample short-read metagenomes

Next, we analyzed fairy versus BWA across different binning algorithms (**Fig. 3A** and **3B**). We focus on **multi-sample, short-read** binning. The mean percentage of recovered bins (*>* 50% complete; *<* 5% contaminated) as a percentage of BWA’s # of bins is 88.6%, 98.5%, 99.1%, and 83.8% for MaxBin2, MetaBAT2, MetaBinner, and VAMB respectively. MaxBin2 and VAMB performed noticeably worse when using fairy. For VAMB, our single-sample assembly with multi-sample coverage approach differs from the “multisplit” approach that VAMB suggests using, perhaps impacting performance. MaxBin2 failed to complete for the chicken caecum dataset where fairy outperformed BWA on all other binners (102%, 102%, and 101% recovered compared to BWA for MetaBAT2, MetaBinner, and VAMB).

**Fig. 3.**
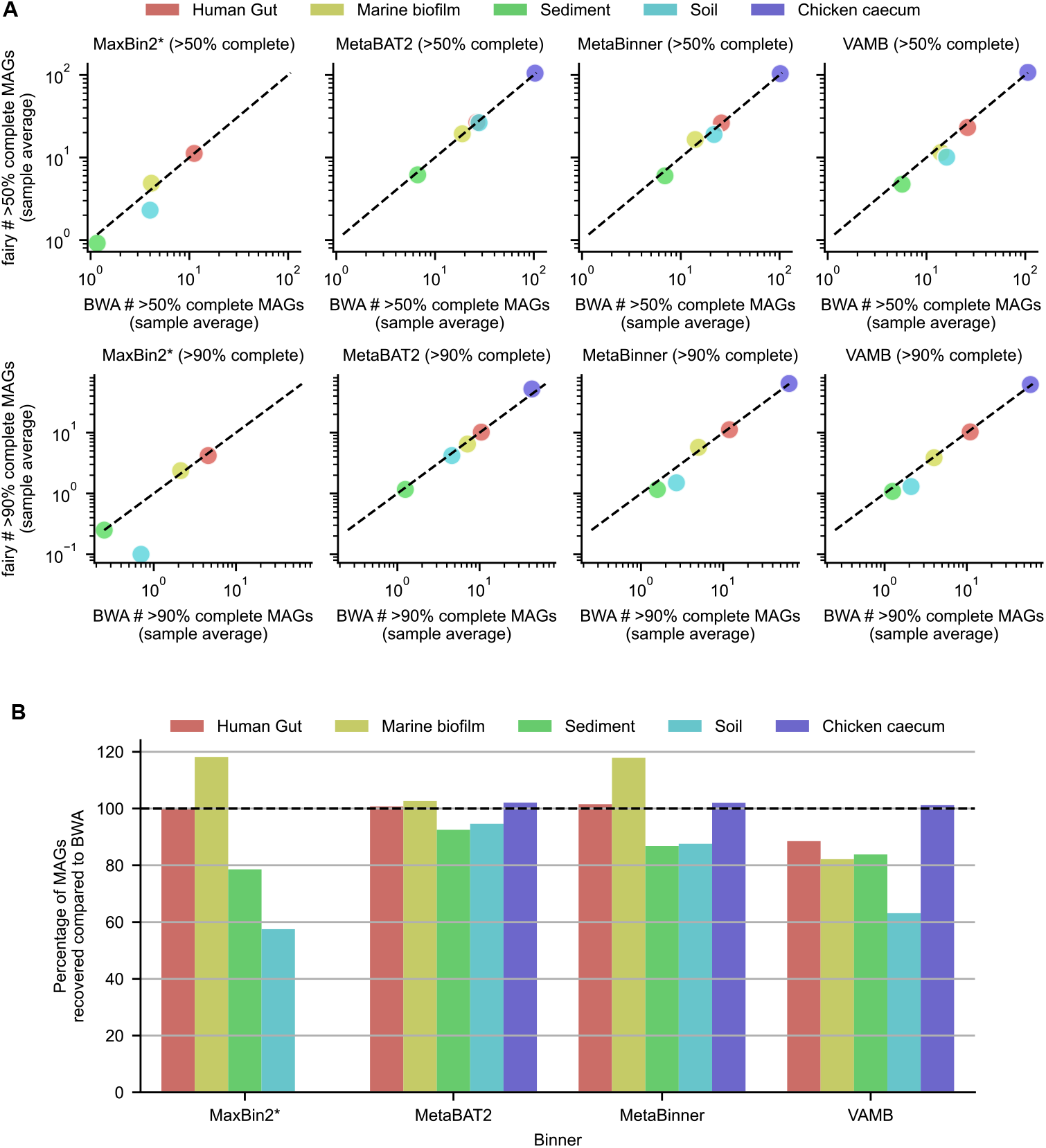
**A**. Fairy’s concordance with BWA for multi-sample, short-read binning. The average number of bins (per sample) with *>* 50 or 90% completeness and *<* 5% contamination is displayed. *MaxBin2 could not run for the chicken caecum dataset due to a software error. **B**. Percentage of bins in (A) with *>* 50% completeness and *<* 5% contamination obtained with fairy relative to BWA over four different binners. This was calculated as 100 (# of fairy bins)*/*(# of BWA bins). Higher than 100% indicates superior performance to BWA. Complete results are available in Supplementary Table 1.

On our host-associated datasets (human gut and chicken caecum), fairy produces more *>* 50% complete bins than BWA for all binners except VAMB. Fairy particularly excelled for high-quality bins (*>* 90% complete and *<* 5% contaminated) on the caecum dataset; fairy recovered 120% of BWA’s bins on the caecum dataset using MetaBAT2, a 20% increase in recovery.

### 3.4 When to use fairy versus alignment

Our results suggest that fairy performs well for multi-sample binning and especially for host-associated metagenomes. We outline two situations below where our results suggest that fairy can not yet replace read alignment.

#### 3.4.1 Caveat 1: fairy is not as good as alignment for single-sample binning

For *single-sample* coverage with short reads, the performance of fairy is noticeably worse than BWA (**Fig. 2**). Using MetaBAT2, fairy recovers only 60% of BWA’s highquality (*>* 90% completeness and *<* 5% contamination) human gut bins for singlesample binning, but 97% of BWA’s high-quality bins for multi-sample binning. We do not recommend fairy for single-sample binning – single-sample coverage calculation is not a bottleneck anyway.

#### 3.4.2 Caveat 2: fairy is usable with nanopore long-reads, but not high-fidelity reads

For nanopore long-reads, our MetaBAT2 results show that fairy is competitive with minimap2’s alignment coverage (**Fig. 2**). However, fairy’s results are markedly worse than minimap2 when using PacBio HiFi reads. We found that this is because HiFi assemblers assemble similar strains instead of collapsing them, and our k-mer-based approach is not suitable for calculating coverage for extremely similar strains.

It may however be possible to tune fairy’s ANI thresholds for strain-level HiFi binning. Although PacBio HiFi metagenomic samples are still uncommon relative to short reads, we believe that strain-level binning is an interesting avenue to explore for future work. Nevertheless, we do not recommend fairy in its current version for PacBio HiFi MAG recovery.

### 3.5 Case study: fairy recovers comparable Asgard archaea genomes in sediment

The study of Asgard archaea (or Asgardarchaeota) has served an important role in the study of eukaryogenesis [41]. Asgard archaea genomes were first recovered through metagenomics [42], and metagenomics continues to be important in their study [28]. In **Fig. 4**, we investigate the quality of Asgard archaea MAGs recovered from the sediment dataset (also used in **Fig. 2, 3**). Sediment metagenomes are known to be complex [43], so this presents an interesting challenge for fairy.

**Fig. 4.**
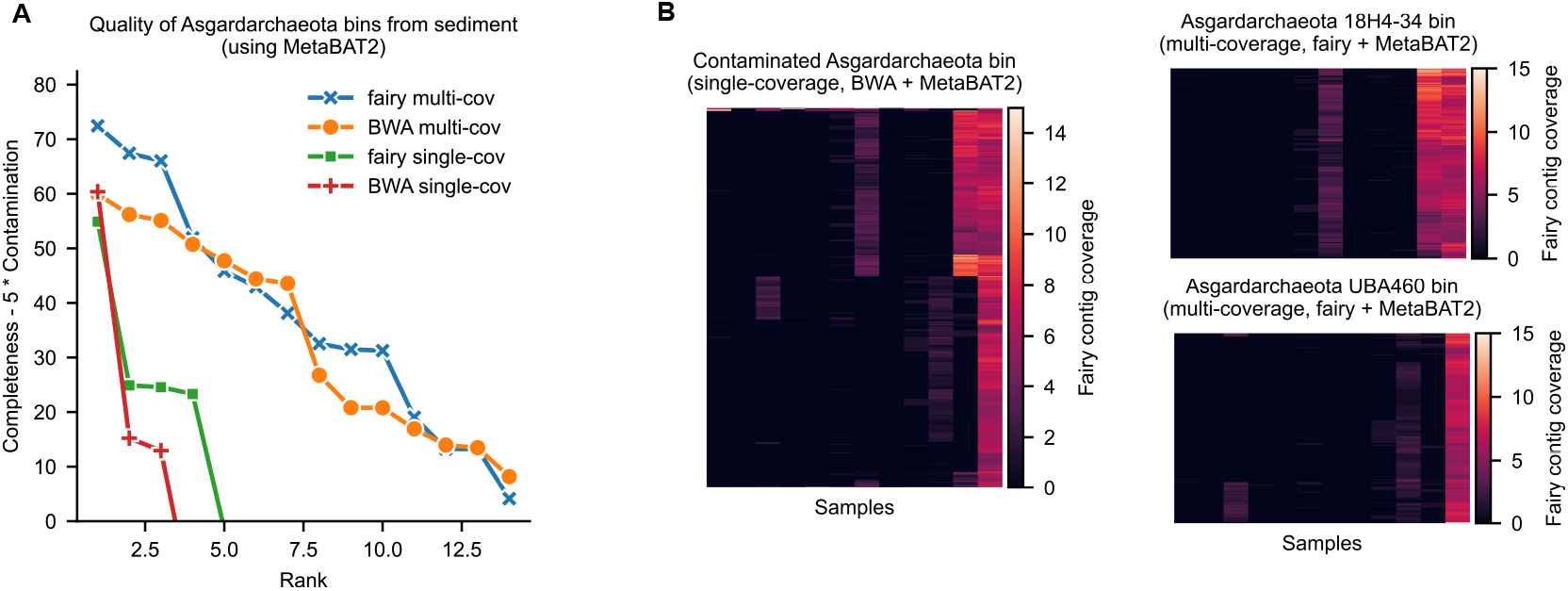
Investigating Asgard archaea from sediment metagenomes using MetaBAT2 with fairy and BWA. **A**. Multi-sample binning, whether with fairy or BWA, recovers much higher quality Asgard archaea bins. **B**. Coverage-contig heatmaps for specific bins. Left: single-sample binning produces a contaminated bin with two Asgard archaea (BWA’s bin pictured). Coverage patterns (fairy’s coverage) show clear contamination. Right: multi-sample binning disentangle the two genomes and correctly generates two bins (fairy’s bins pictured).

For MetaBAT2 bins from the sediment metagenomes, we compared single-sample versus multi-sample binning and found that multi-sample binning produces much higher quality Asgard bins (**Fig. 4A**). Fairy is even slightly better than BWA when ranking the bins under the Completeness −5× Contamination metric.

We visualized an explicit example of where single-sample coverage binning failed in **Fig. 4B**. In the sample DRR310882, both fairy and BWA (with MetaBAT2) produced a contaminated bin with two Asgard archaea. BWA’s bin (visualized in **Fig. 4B**, left) had a 55% contamination estimate. The two erroneously binned Asgard genomes have similar coverage in this sample and could not be separated with k-mer frequencies. However, multi-sample coverage information disentangles the two genomes, and both fairy and BWA do so successfully. Fairy’s resulting multi-sample bins (**Fig. 4B**, right), with genus assignments 18H4-34 and UBA460 using GTDB-Tk [44], had only 2.66% and 4.25% contamination respectively.

## 4 Discussion

### 4.1 Fairy’s role in MAG recovery pipelines

Fairy is orders of magnitude faster than read alignment and solves a key computational bottleneck. The speed increase is not controversial. In some cases, fairy may give better results (e.g. the biofilm and chicken caecum datasets in **Fig. 2**), but there may be a slight sensitivity loss compared to read alignment on complex metagenomes such as sediment or soil. However, we show that when all-to-all alignment is not feasible, multi-sample binning with fairy should always be preferred over single-sample read alignment, thus filling an important niche for large-sample projects.

Furthermore, we envision fairy as complementary to all-to-all read alignment. Users can first use fairy to obtain a set of good-quality multi-sample bins and immediately analyze their data while they wait days/weeks for their all-to-all read alignments finish.

### 4.2 For default, non-tuned binning parameters, fairy’s accuracy is encouraging

Binning accuracy depends on both the binner *and* the coverage calculation method. We used standard pipelines for obtaining read-alignment coverage, but binning algorithms are designed with these pipelines in mind and tuned specifically for read-alignment coverages. This disadvantages fairy, whose coverage statistics likely have peculiarities that binners have not been tuned for. In the future, binners that are designed with kmer-based coverage in mind should be more performant relative to read alignment than our current results suggest. We could have, in theory, tested out different parameters to optimize the binners for each type of data, but we decided that this should be investigated in future work instead.

## 5 Conclusion

In this paper, we provide a new k-mer-based coverage calculation method for metagenomic binning called fairy. Fairy is magnitudes faster than read alignment and enables multi-sample binning on much larger datasets than before while costing only a bit of sensitivity.

## Availability of Data and Materials

The latest version of fairy is available at https://github.com/bluenote-1577/fairy. The benchmarked version 0.5.1 is available at https://github.com/bluenote-1577/fairy/tree/4d8e8eba5620629716b538f3e866fc4605b17ede. All read accessions used for benchmarking are available in Supplementary Table 2. Scripts for recreating the figures are available at https://github.com/bluenote-1577/fairy-test.

## Contributions

J.S. devised the study, ran the experiments, and wrote the software. Y.W.Y supervised the study. Both authors wrote and edited the manuscript.

## A Containment ANI and coverage calculation

### A.1 FracMinHash for k-mer sampling

FracMinHash [23] is a method for systematically selecting a subset of k-mers from a larger set of k-mers (i.e. subsampling). All k-mers used in fairy are obtained after selection by FracMinHash. We describe FracMinHash below.

Let *h* : Σ^*k*^ →{0, …, 2^64^ −1} be a hash function from k-mers to 64-bit integers. Let *c* be a positive number. Given a set of k-mers *X*, we obtain a subset

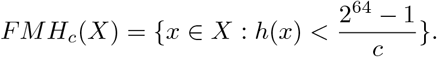

Intuitively, FracMinHash returns a set of k-mers approximately *c* times smaller than the original set, assuming a reasonably uniform hash function. The key reason we use FracMinHash is that k-mers are *consistently* subsampled across different sequences – given a fixed hash function, if a k-mer is sampled on a contig, the same k-mer will be sampled on the read (because its hash value is the same).

In practice, fairy uses minimap2’s [37] hash function. Fairy uses *c* = 50 by default and *k* = 31.

### A.2 Containment ANI

Let *A* be a contig’s k-mers after applying FracMinHash. Let *B* be a metagenomic sample’s (i.e. a set of reads) collection of k-mers after applying FracMinHash (over *all* reads). The naive containment ANI is defined as

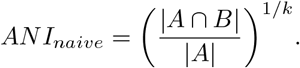

The containment ANI generalizes the standard ANI for genome-to-genome comparisons [45] under a simple statistical model. The naive containment ANI is not accurate when the coverage of the contig is low within the sample [46]. To deal with this, sylph [21] introduced a statistical procedure to estimate containment ANI accurately under low coverage.

Briefly, let *N*_*a*_ be the number of k-mers in *A* (the contig) with multiplicity/count *a* in *B* (the sample). Under stochastic sequencing assumptions, we can estimate the *effective coverage* as 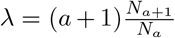 [21, 47] for any *a >* 0; *a* is chosen to correspond to the largest value of *N*_*a*_. Then, fairy’s containment ANI can be estimated as

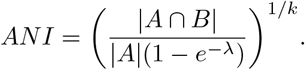

In practice, fairy only estimates *λ* when the median k-mer coverage in the sample is ≤ 3, otherwise we use the naive formula. Additional implementation details are given in the Methods section of sylph’s manuscript [21].

### Coverage calculation

Given a set of multiplicities for the k-mers in *A* (the contig) in the metagenomic sample, we calculate coverage as follows. Let *M* be the median k-mer multiplicity.

- If *M* ≤ 3: fairy outputs the *λ* estimate (discussed previously).
- If 4 ≤ *M*≤ 15: fairy uses a robust mean as follows. Let *Z* ∼ *Pois*(*M*) be a Poisson random variable with mean *M*, the median k-mer multiplicity. We discard all k-mer multiplicities *α* with *P* (*Z > α*) *<* 10^−10^, and then output a robust mean. This is done to discard long-tailed outliers due to repetitive and shared k-mers.
- If *M >* 15, fairy outputs the median k-mer multiplicity *M* .

The intuition for the above procedure is that for small *M*, a statistical method is crucial because means and medians do not give enough resolution. For moderate *M*, means give better resolution than medians but require thresholding to ensure robustness to outliers.

## References

[1] Quince, C., Walker, A.W., Simpson, J.T., Loman, N.J., Segata, N.: Shotgun metagenomics, from sampling to analysis. Nature Biotechnology 35(9), 833–844 (2017) 10.1038/nbt.3935

[2] Kang, D.D., Li, F., Kirton, E., Thomas, A., Egan, R., An, H., Wang, Z.: MetaBAT 2: An adaptive binning algorithm for robust and efficient genome reconstruction from metagenome assemblies. PeerJ 7 (2019) 10.7717/peerj.7359

[3] Wu, Y.-W., Simmons, B.A., Singer, S.W.: MaxBin 2.0: An automated binning algorithm to recover genomes from multiple metagenomic datasets. Bioinformatics 32(4), 605–607 (2016) 10.1093/bioinformatics/btv638

[4] Nissen, J.N., Johansen, J., Allesøe, R.L., Sønderby, C.K., Armenteros, J.J.A., Grønbech, C.H., Jensen, L.J., Nielsen, H.B., Petersen, T.N., Winther, O., Rasmussen, S.: Improved metagenome binning and assembly using deep variational autoencoders. Nature Biotechnology 39(5), 555–560 (2021) 10.1038/s41587-020-00777-4

[5] Wang, Z., Huang, P., You, R., Sun, F., Zhu, S.: MetaBinner: A high-performance and stand-alone ensemble binning method to recover individual genomes from complex microbial communities. Genome Biology 24(1), 1 (2023) 10.1186/s13059-022-02832-6

[6] Pavia, M.J., Chede, A., Wu, Z., Cadillo-Quiroz, H., Zhu, Q.: BinaRena: A dedicated interactive platform for human-guided exploration and binning of metagenomes. Microbiome 11(1), 186 (2023) 10.1186/s40168-023-01625-8

[7] Dick, G.J., Andersson, A.F., Baker, B.J., Simmons, S.L., Thomas, B.C., Yelton, A.P., Banfield, J.F.: Community-wide analysis of microbial genome sequence signatures. Genome Biology 10(8), 85 (2009) 10.1186/gb-2009-10-8-r85

[8] Alneberg, J., Bjarnason, B.S., de Bruijn, I., Schirmer, M., Quick, J., Ijaz, U.Z., Lahti, L., Loman, N.J., Andersson, A.F., Quince, C.: Binning metagenomic contigs by coverage and composition. Nature Methods 11(11), 1144–1146 (2014) 10.1038/nmeth.3103

[9] Chklovski, A., Parks, D.H., Woodcroft, B.J., Tyson, G.W.: CheckM2: A rapid, scalable and accurate tool for assessing microbial genome quality using machine learning. Nature Methods 20(8), 1203–1212 (2023) 10.1038/s41592-023-01940-w

[10] Mattock, J., Watson, M.: A comparison of single-coverage and multi-coverage metagenomic binning reveals extensive hidden contamination. Nature Methods, 1–4 (2023) 10.1038/s41592-023-01934-8

[11] Li, H.: Aligning sequence reads, clone sequences and assembly contigs with BWA-MEM (1303.3997) (2013) 10.48550/arXiv.1303.3997 arxiv:1303.3997 [q-bio]

[12] Langmead, B., Salzberg, S.L.: Fast gapped-read alignment with Bowtie 2. Nature methods 9(4), 357–359 (2012) 10.1038/nmeth.1923

[13] Olm, M.R., Brown, C.T., Brooks, B., Banfield, J.F.: dRep: A tool for fast and accurate genomic comparisons that enables improved genome recovery from metagenomes through de-replication. The ISME Journal 11(12), 2864–2868 (2017) 10.1038/ismej.2017.126

[14] Pasolli, E., Asnicar, F., Manara, S., Zolfo, M., Karcher, N., Armanini, F., Beghini, F., Manghi, P., Tett, A., Ghensi, P., Collado, M.C., Rice, B.L., DuLong, C., Morgan, X.C., Golden, C.D., Quince, C., Huttenhower, C., Segata, N.: Extensive Unexplored Human Microbiome Diversity Revealed by Over 150,000 Genomes from Metagenomes Spanning Age, Geography, and Lifestyle. Cell 176(3), 649–66220 (2019) 10.1016/j.cell.2019.01.001

[15] Bray, N.L., Pimentel, H., Melsted, P., Pachter, L.: Near-optimal probabilistic RNA-seq quantification. Nature Biotechnology 34(5), 525–527 (2016) 10.1038/nbt.3519

[16] Patro, R., Duggal, G., Love, M.I., Irizarry, R.A., Kingsford, C.: Salmon provides fast and bias-aware quantification of transcript expression. Nature Methods 14(4), 417–419 (2017) 10.1038/nmeth.4197

[17] Alneberg, J., Bennke, C., Beier, S., Bunse, C., Quince, C., Ininbergs, K., Riemann, L., Ekman, M., Jürgens, K., Labrenz, M., Pinhassi, J., Andersson, A.F.: Ecosystem-wide metagenomic binning enables prediction of ecological niches from genomes. Communications Biology 3(1), 1–10 (2020) 10.1038/s42003-020-0856-x

[18] Zorrilla, F., Buric, F., Patil, K.R., Zelezniak, A.: metaGEM: Reconstruction of genome scale metabolic models directly from metagenomes. Nucleic Acids Research 49(21), 126 (2021) 10.1093/nar/gkab815

[19] Quince, C., Nurk, S., Raguideau, S., James, R., Soyer, O.S., Summers, J.K., Limasset, A., Eren, A.M., Chikhi, R., Darling, A.E.: STRONG: Metagenomics strain resolution on assembly graphs. Genome Biology 22(1), 214 (2021) 10.1186/s13059-021-02419-7

[20] Krivonosova, K., Gorshkov, Y., Nurk, S.: Estimating Differential Abundance Profiles for Metagenomic Series Binning. https://bioinformaticsinstitute.ru/sites/default/files/4krivonosova150918.pdf

[21] Shaw, J., Yu, Y.W.: Metagenome profiling and containment estimation through abundance-corrected k-mer sketching with sylph. bioRxiv, 2023–1120567879 (2023) 10.1101/2023.11.20.567879

[22] Yorukoglu, D., Yu, Y.W., Peng, J., Berger, B.: Compressive mapping for nextgeneration sequencing. Nature Biotechnology 34(4), 374–376 (2016) 10.1038/nbt.3511

[23] Irber, L., Brooks, P.T., Reiter, T., Pierce-Ward, N.T., Hera, M.R., Koslicki, D., Brown, C.T.: Lightweight compositional analysis of metagenomes with FracMin-Hash and minimum metagenome covers, 2022–0111475838 (2022) 10.1101/2022.01.11.475838 [New Results]. Chap. New Results

[24] Jain, C., Rodriguez-R, L.M., Phillippy, A.M., Konstantinidis, K.T., Aluru, S.: High throughput ANI analysis of 90K prokaryotic genomes reveals clear species boundaries. Nature Communications 9(1), 5114 (2018) 10.1038/s41467-018-07641-9

[25] Feng, X., Cheng, H., Portik, D., Li, H.: Metagenome assembly of high-fidelity long reads with hifiasm-meta. Nature Methods 19(6), 671–674 (2022) 10.1038/s41592-022-01478-3

[26] Glendinning, L., Stewart, R.D., Pallen, M.J., Watson, K.A., Watson, M.: Assembly of hundreds of novel bacterial genomes from the chicken caecum. Genome Biology 21(1), 34 (2020) 10.1186/s13059-020-1947-1

[27] Olm, M.R., Butterfield, C.N., Copeland, A., Boles, T.C., Thomas, B.C., Banfield, J.F.: The Source and Evolutionary History of a Microbial Contaminant Identified Through Soil Metagenomic Analysis. mBio 8(1), 10–11280196916 (2017) 10.1128/mbio.01969-16

[28] Medvedeva, S., Sun, J., Yutin, N., Koonin, E.V., Nunoura, T., Rinke, C., Krupovic, M.: Three families of Asgard archaeal viruses identified in metagenome-assembled genomes. Nature Microbiology 7(7), 962–973 (2022) 10.1038/s41564-022-01144-6

[29] Zhang, W., Ding, W., Li, Y.-X., Tam, C., Bougouffa, S., Wang, R., Pei, B., Chiang, H., Leung, P., Lu, Y., Sun, J., Fu, H., Bajic, V.B., Liu, H., Webster, N.S., Qian, P.-Y.: Marine biofilms constitute a bank of hidden microbial diversity and functional potential. Nature Communications 10, 517 (2019) 10.1038/s41467-019-08463-z

[30] Sereika, M., Petriglieri, F., Jensen, T.B.N., Sannikov, A., Hoppe, M., Nielsen, P.H., Marshall, I.P.G., Schramm, A., Albertsen, M.: Closed genomes uncover a saltwater species of Candidatus Electronema and shed new light on the boundary between marine and freshwater cable bacteria. The ISME Journal 17(4), 561–569 (2023) 10.1038/s41396-023-01372-6

[31] Gounot, J.-S., Chia, M., Bertrand, D., Saw, W.-Y., Ravikrishnan, A., Low, A., Ding, Y., Ng, A.H.Q., Tan, L.W.L., Teo, Y.-Y., Seedorf, H., Nagarajan, N.: Genome-centric analysis of short and long read metagenomes reveals uncharacterized microbiome diversity in Southeast Asians. Nature Communications 13(1), 6044 (2022) 10.1038/s41467-022-33782-z

[32] Sidhu, C., Kirstein, I.V., Meunier, C.L., Rick, J., Fofonova, V., Wiltshire, K.H., Steinke, N., Vidal-Melgosa, S., Hehemann, J.-H., Huettel, B., Schweder, T., Fuchs, B.M., Amann, R.I., Teeling, H.: Dissolved storage glycans shaped the community composition of abundant bacterioplankton clades during a North Sea spring phytoplankton bloom. Microbiome 11(1), 77 (2023) 10.1186/s40168-023-01517-x

[33] Uritskiy, G.V., DiRuggiero, J., Taylor, J.: MetaWRAP—a flexible pipeline for genome-resolved metagenomic data analysis. Microbiome 6(1), 158 (2018) 10.1186/s40168-018-0541-1

[34] Kieser, S., Brown, J., Zdobnov, E.M., Trajkovski, M., McCue, L.A.: ATLAS: A Snakemake workflow for assembly, annotation, and genomic binning of metagenome sequence data. BMC Bioinformatics 21(1), 257 (2020) 10.1186/s12859-020-03585-4

[35] Aroney, S.T.N., Newell, R.J.P., Nissen, J., Camargo, A.P., Tyson, G.W., Woodcroft, B.J.: CoverM: Read Coverage Calculator for Metagenomics. Zenodo (2024). 10.5281/zenodo.10531254

[36] Kolmogorov, M., Bickhart, D.M., Behsaz, B., Gurevich, A., Rayko, M., Shin, S.B., Kuhn, K., Yuan, J., Polevikov, E., Smith, T.P.L., Pevzner, P.A.: metaFlye: Scalable long-read metagenome assembly using repeat graphs. Nature Methods 17(11), 1103–1110 (2020) 10.1038/s41592-020-00971-x

[37] Li, H.: Minimap2: Pairwise alignment for nucleotide sequences. Bioinformatics 34(18), 3094–3100 (2018) 10.1093/bioinformatics/bty191

[38] Benoit, G., Raguideau, S., James, R., Phillippy, A.M., Chikhi, R., Quince, C.: High-quality metagenome assembly from long accurate reads with metaMDBG. Nature Biotechnology, 1–6 (2024) 10.1038/s41587-023-01983-6

[39] Pan, S., Zhao, X.-M., Coelho, L.P.: SemiBin2: Self-supervised contrastive learning leads to better MAGs for short- and long-read sequencing. Bioinformatics 39(Supplement 1), 21–29 (2023) 10.1093/bioinformatics/btad209

[40] Liu, C.-C., Dong, S.-S., Chen, J.-B., Wang, C., Ning, P., Guo, Y., Yang, T.-L.: MetaDecoder: A novel method for clustering metagenomic contigs. Microbiome 10(1), 46 (2022) 10.1186/s40168-022-01237-8

[41] Eme, L., Tamarit, D., Caceres, E.F., Stairs, C.W., De Anda, V., Schön, M.E., Seitz, K.W., Dombrowski, N., Lewis, W.H., Homa, F., Saw, J.H., Lombard, J., Nunoura, T., Li, W.-J., Hua, Z.-S., Chen, L.-X., Banfield, J.F., John, E.S., Reysenbach, A.-L., Stott, M.B., Schramm, A., Kjeldsen, K.U., Teske, A.P., Baker, B.J., Ettema, T.J.G.: Inference and reconstruction of the heimdallarchaeial ancestry of eukaryotes. Nature 618(7967), 992–999 (2023) 10.1038/s41586-023-06186-2

[42] Spang, A., Saw, J.H., Jørgensen, S.L., Zaremba-Niedzwiedzka, K., Martijn, J., Lind, A.E., van Eijk, R., Schleper, C., Guy, L., Ettema, T.J.G.: Complex archaea that bridge the gap between prokaryotes and eukaryotes. Nature 521(7551), 173–179 (2015) 10.1038/nature14447

[43] Chevrette, M.G., Bratburd, J.R., Currie, C.R., Stubbendieck, R.M.: Experimental Microbiomes: Models Not to Scale. mSystems 4(4), 10–11280017519 (2019) 10.1128/msystems.00175-19

[44] Chaumeil, P.-A., Mussig, A.J., Hugenholtz, P., Parks, D.H.: GTDB-Tk: A toolkit to classify genomes with the Genome Taxonomy Database. Bioinformatics 36(6), 1925–1927 (2020) 10.1093/bioinformatics/btz848

[45] Richter, M., Rosselló-Móra, R.: Shifting the genomic gold standard for the prokaryotic species definition. Proceedings of the National Academy of Sciences 106(45), 19126–19131 (2009) 10.1073/pnas.0906412106

[46] Ondov, B.D., Treangen, T.J., Melsted, P., Mallonee, A.B., Bergman, N.H., Koren, S., Phillippy, A.M.: Mash: Fast genome and metagenome distance estimation using MinHash. Genome Biology 17(1), 132 (2016) 10.1186/s13059-016-0997-x

[47] Sarmashghi, S., Bohmann, K., P. Gilbert, M.T., Bafna, V., Mirarab, S.: Skmer: Assembly-free and alignment-free sample identification using genome skims. Genome Biology 20(1), 34 (2019) 10.1186/s13059-019-1632-4

